# Extracellular electrophysiological effects on MEA recordings of living mycelium following in-vivo polydopamine polymerisation

**DOI:** 10.1101/2025.09.11.675229

**Authors:** Davin Browner, Andrew Adamatzky

## Abstract

Fungi produce multiple melanins found in nature, including the eumelanins (black or dark brown), pheomelanins (yellow or red) and the highly diverse group of allomelanins. The fungal melanins provide protection against UV radiation, oxidative stress, desiccation, host immune evasion and antifungal drug resistance. As a result, research on melanin in fungi is wide ranging including development of anti-fungals and surface interfacing bioelectronics. In-vivo polymerisation of compounds with electrical or ionic conductivity can provide a useful method to interface with the electrophysiology of biological organisms. However, the impact of such interventions is unknown where anti-fungal activity is present and interferes with the target signaling in the organism. Melanin compounds are promising for achieving bioelectronic interfaces with mycelium. However, the identification of the role of melanin in blocking calcium flux in airway epithelial cells and phagosomes underscores the significance of investigating its effects in more detail. Additionally, synthetic analogs such as polydopamine are thought to exhibit similar metal ion chelation properties, supporting the biochemical parallels between natural and artificial melanins. In this paper, we investigate the polymerisation of synthetic melanins in mycelium from the intracellular environment and onto microelectrodes in terms of extracellular electrophysiology. At concentrations of 5 mM and 10 mM extracellular electrophysiological activity is significantly inhibited suggesting a putative link to reduction in Ca^2+^ efflux and antifungal properties. The dispersed mycelial cultures were estimated to comprise of a total of 200 spiking units across triplicates (T1= 117, T2=40, T3=43). The triplicates had a combined mean trough-to-peak time of 1.72 ± 0.07 ms. Selection of electrophysiology interfacing polymers should be carefully orchestrated to avoid impacting baseline physiology of fungi.

## Introduction

Fungi are known to produce multiple types of melanin, primarily DOPA-melanin (eumelanin), DHN-melanin, and in some cases pyomelanin. Melanin is formed by the oxidative polymerization of phenolic or indolic compounds (like L-DOPA, catechol, or 1,8-DHN). The structure and properties of the melanin depend on the precursor. For example, 1,8-dihydroxynaphthalene (DHN) leads to DHN-melanin, while L-DOPA leads to DOPA-melanin [1]. Despite different biosynthetic pathways, melanins share traits like hue, UV protection, free radical scavenging and metal ion binding. Melanin plays multiple roles in fungi: protection against UV radiation, oxidative stress, desiccation, host immune evasion, and antifungal drug resistance. Fungi produce a wide range of melanins found in nature, including eumelanins (black or dark brown), pheomelanins (yellow or red), and the highly diverse group of allomelanins. The latter includes soluble pyomelanins and melanins derived from 1,8-dihydroxynaphthalene (DHN) compounds [1]. These melanins confer significant protection against environmental and host-derived stresses—such as UV radiation, oxidative damage, enzymatic attack, and antimicrobial agents—thereby enhancing fungal survival and promoting pathogenicity. Beyond passive protection, melanin can actively contribute to virulence by aiding in tissue invasion and suppressing host immune responses, facilitating the establishment and persistence of infection.

Research on melanin Ca^2+^ efflux blocking in mycelium has emerged as an important experimental modality due to its implications for fungal pathogenicity, immune evasion, and potential biomedical applications. Fungal melanins, including dihydroxynaphthalene (DHN)-melanin and DOPA-melanin, have been extensively studied for their roles in protecting fungi from environmental stresses and host immune responses [2, 3]. Research has explored melanin’s involvement in fungal virulence, particularly in species such as *Aspergillus fumigatus* and *Cryptococcus neoformans*, where melanin modulates host-pathogen interactions and contributes to resistance against antifungal agents [4, 5].

Despite advances in understanding melanin biosynthesis and its protective functions, the precise biochemical and biophysical interactions underlying Ca2+ blocking remain incompletely characterised [6, 7, 8, 9]. The extent to which different melanin types contribute to calcium sequestration and immune modulation varies, with some studies emphasizing DHN-melanin’s role in inhibiting calcium-calmodulin signaling, while others highlight the structural and functional diversity of melanins influencing these processes [10, 11, 12].

A number of studies quantitatively demonstrated fungal melanin’s capacity to sequester Ca^2+^ [4, 6, 9]. Melanin’s calcium binding was shown to limit calcium availability for critical fungal processes such as capsule assembly, impacting fungal physiology and host interactions [6, 8]. Modified melanins with enhanced solubility or altered structure exhibited varied calcium binding affinities, suggesting potential for tuning melanin’s biochemical functions [13]. The modificaiton of these biochemical functions may be relevant to fungal electrophysiology and bioelectronics [14, 15]. However, studies reported that melanin-mediated calcium sequestration disrupts intracellular calcium flux and downstream signaling pathways, notably calcium-calmodulin signaling essential for immune responses and fungal survival [4, 6, 7]. Melanin’s interference with calcium signaling has also been linked to inhibition of phagosome maturation and autophagy-related pathways, impairing host immune cell functions [6, 8]. Some studies implied that melanin’s calcium blocking indirectly modulates fungal metabolic pathways and drug susceptibility through altered signaling [7]. As a result, use of synthetic melanins for fungal electrophysiology requires careful consideration of impacts on ion channel activity including Ca^2+^ efflux.

The differences in Ca^2+^ efflux inhibition between fungal melanins and synthetic melanins such as polydopamine remains underexplored. Fungal melanin is a heterogeneous polymeric pigment synthesized via DHN or DOPA pathways and exhibits significant Ca^2+^ chelation and ion channel modulation. In comparison, polydopamine as a synthetic melanin mimic with similar physicochemical properties. Calcium ion sequestration is a critical modulatory mechanism affecting fungal-host interactions [16, 17, 5].

Multiple studies employed advanced techniques such as solid-state NMR and biochemical assays to characterize fungal melanins and polydopamine, revealing heterogeneous polymeric structures with metal-binding sites relevant to calcium interaction [18, 5]. Structural analyses highlighted differences between natural fungal melanins (e.g., DHN-melanin, DOPA-melanin) and synthetic polydopamine, yet confirmed functional similarities in calcium chelation [18]. The spatial arrangement of melanin within fungal cell walls and its association with other cell wall components were emphasized as critical for calcium binding and biological activity [8]. Studies demonstrated that melanin’s calcium blocking leads to significant immunomodulatory effects, including suppression of chemokine secretion, reduced neutrophil recruitment and inhibition of phagocytosis [4, 6]. Synthetic melanin analogs like polydopamine were also noted for their potential to modulate immune responses via calcium chelation [18]. Polydopamine exhibits metal ion chelation and calcium-binding properties analogous to natural fungal melanins, enabling its use as a synthetic model for studying melanin-Ca^2+^ interactions. Its structural tailoring and radical scavenging capabilities facilitate exploration of melanin’s functional roles in biomedical applications, including modulation of calcium flux in biological systems [17]. These biochemical qualities make it suitable for investigation of bioelectronic, protonic and Ca^2+^ interfaces with mycelium. However, the influence of melanin on Ca^2+^ channels and transporters remains underexplored, representing a significant gap in understanding the full scope of melanin-mediated calcium flux modulation in fungal cells and host interactions. The degree to which melanin modulates calcium signaling varies, with some effects not fully elucidated. Variation in host cell models (human airway epithelium, macrophages, amoebae), fungal species, and melanin type may explain differences in calcium signaling impact. Experimental approaches vary in resolution and scope. As a result, studies could be aided by high temporal resolution investigation of extracellular calcium flux upon exposure to melanins utilising standardised electrophysiological tools such as microelectrode arrays.

Modulation of calcium ion dynamics within fungal cells, particularly through sequestration and blocking of Ca^2+^ flux could also prove to be useful for identifying the physiological origin of fungal extracellular electrophysiology. Early research on fungal electrical signaling suggested the role of proton pumps (H^+^) and various ion channels (Ca^2+^, Cl^-^) [19, 20] utilising patch clamp intracellular recordings methods. However, in extracellular studies specific mappings from active or passive ion channels to discrete or continuous type signals are not currently well understood at the wide spectral bandwidths typically associated with electrophysiological studies (0.01-6000 Hz) and resulting temporal dynamics.

In this paper, we investigate the morphological growth of Basidiomycete mycelium in liquid cultures with polydopamine (10 mM and 5 mM solutions) to assess the effects on extracellular electrophysiology via microelectrode array recordings. At the surveyed concentrations polydopamine appears to block electrophysiological discrete unit spikes (150-3000 Hz) at 5 mM and 10 mM suggesting a putative link to Ca^2+^ efflux. However, different synthesis pathways may be required to utilise polydopamine as a bioelectrical interface if the target electrophysiological processes involve calcium efflux. Melanins may be engineered to produce different effects on Ca^2+^ efflux with impact on biocomputing, sensing and treatment of fungal diseases. Fungal melanins and close synthetic analogs remain as a promising biochemical interface for studies of pathogenicity and bioelectronics via the endogenous electrophysiology of mycelium.

## Methods

### Liquid culture cultivation

*Hericium erinaceus* cultures were sampled from stocks and grown in flasks using sterilised malt liquid culture media (1.5 %). Aeration of the growing cultures was achieved using two syringe Filters (13mm diameter and 0.22 *µ*m filter size). Constant stirring was utilised during growth to avoid formation of exopolymeric substances. Cultures were grown for 2 days using this setup before introduction of polydopamine.

### In-vivo polymerisation of polydopamine in liquid cultures

Polydopamine was synthesised using dopamine hydrochloride (Sigma, U.S.). The dopamine hydrochloride was prepared as 10 mM and 5 mM solutions in deionised water. 1 ml of dispersed liquid culture solution was then transferred to the polydopamine solution and left to grow overnight. The polymerisation resulted transition from a teal coloured liquid to a dark brown liquid covering mycelium overnight in the 10 mM cultures. In the 5 mM cultures the solution transitioned from clear to teal over the same period. This indicated that polymerisation had occured in both cases with different densities. We confirmed intracellular polymerisation using environmental scanning electron microscopy (ESEM). The FEI Quanta 650 FEG scanning electron microscope (SEM) (FEI Company, U.S.) was used to image the samples in a hydrated state and produce micrographs. Selected liquid cultures were placed on a stainless steel stub on the Peltier stage that was set to cool to 2^°^C. At this temperature, the dew point of water was lowered below 10 Torr. A drop of distilled water was placed in each of the four depressions in the Peltier substage to promote rapid increase in humidity. The chamber was pumped to 7.5 Torr. The set pressure was then reduced in 0.2 Torr steps to 6.7 Torr. The surface of the water drop was observed, and the water evaporated slowly by reducing the pressure to expose the fully hydrated samples. A repeated humidity modification was used for each sample dropping to 90 % and remaining at this humidity level for the duration of imaging.

### Micro-electrode arrays (MEAs)

The microelectrode array (MEA) was a custom printed circuit board (PCB) with 10U” hard gold coating. The MEA had 64 channels including a reference electrode. The single ended circular electrodes in the MEA are arranged in a rectangular grid array with each electrode having a radius of 25 *µ*m and a vertical and horizontal spacing of 700 *µ*m. The exposed electrically conductive surface of the electrodes was equivalent to the diameter at 50 *µ*m each. Recordings were conducted in a shielding box (Tescom, South Korea). The shielding box functioned as a Faraday cage with copper mesh lining and also blocked high frequency noise. With RF shielding effectiveness of up to *>* 80 dB across a wide frequency range, the TC-5910D also acted as a Faraday cage as a result of its conductive copper mesh lining, steel exterior and RF-filtered connectors resulting in a fully enclosed and dark recording environment. The shielding box was placed on an anti-vibration table (Adam Equipment, U.K.) to eliminate mechanical vibration artifacts. Low frequency components and line noise were removed by the bandpass (150-3000 Hz) and common average reference procedures. All samples were rested for 2 hours in-situ to recover from transfer shock.

### Amplification

The Intan RHD2164 (Intan Technologies, U.S.) was used to amplify the signals from the MEA. It is a 64-channel digital electrophysiology interface chip designed for highdensity extracellular recordings. Features 64 low-noise amplifiers with an input-referred noise of 2.4 *µ*V RMS and an input impedance of 1 GΩ // 2 pF. The bandwidth is programmable, with a low cutoff frequency ranging from 0.1 Hz to 500 Hz and a high cutoff frequency from 100 Hz to 20 kHz. The chip includes a 16-bit analog-to-digital converter (ADC) that provides simultaneous sampling of all channels at a maximum sampling rate of 30 kS*/*s per channel. Communication is facilitated through an SPI interface. The MEA was connected via custom housing (connection junctions had a maximum length of 150mm) and routed to the amplifier using an electrode adapter board (Intan Technologies, U.S.). The fungal sample, MEA, and Intan headstages was placed inside the shielding box. The acquisition board controller and computer (which generate digital noise and have high-voltage AC power wiring) were located outside the shielding box. The GND terminal on the Intan headstage was connected to the tissue via a ground reference electrode. The zero-ohm R0 resistor was left in place on the RHD headstage so that REF could be shorted to GND. This resulted in a combined REF/GND electrode that was connected to the microelectrode reference electrode (large triangular gold electrode which removed macroscale electrochemical noise). The fungal sample was electrically isolated from the shielding box and the shielding box was grounded to external electronics. All recordings were conducted in a temperature controlled environment to minimise thermal noise (19^°^C). A butyl rubber lid covered the recording chamber to maintain humidity levels of 90-95 %. Photoelectric noise was negligible due to the dark environment of the interior of the shielding box. The shielding box was locked shut during recordings and was only opened for inspection or changing of samples.

### Data acquisition

The Open Ephys Acquisition Board (OpenEphys, Portugal) was used for data acquisition. It is an open-source interface designed for high-channel-count electrophysiology experiments [21]. In our experiments, a single Intan RHD2164 headstage was connected to the acquisition board via SPI cable. This completed the recording setup and allowed for simultaneously sampling at a per-channel raw recording frequency range of 0.1 to 6500 Hz used in experiments. The combination of Faraday double shielding and RF blocking with a frequency range of 150-3000 Hz for spike detection forms a good foundation for distinguishing between electrophysiological spikes, passive phenomena such as Donnan potentials and environmental noise. Donnan potentials are typically in the 0.1-1 Hz range and therefore are not relevant to the spike detection bandpass utilised here [22]. Similarly mains noise can produce artifacts at 50 Hz. This was below the high pass filter of the bandpass used in sorting and harmonics of this frequency of noise were not detected. All external sensors or sources of noise from the recording chip were configured to the off state (e.g. additional channels or ADC). Grounding and referencing procedures followed guidelines produced by manufacturers [23]. The chamber was sealed from light sources thereby removing the possibility of light-related noise such as the Becquerel effect [24]. Thermal noise artifacts were minimised through the use of a temperature controlled room for all recordings. The recorded potentials are unlikely to be related to physical movement of the mycelium as a result of slow growth dynamics (e.g. electrical artifacts ≤1 Hz). Following acquisition, a digital bandpass filter was used to isolate activity between 150-3000 Hz using a Butterworth second order filter. This frequency range was selected as a result of power spectral density analysis indicating an absence of frequency components ≥3000 Hz and ≤6000 Hz. A common reference electrode was used to reduce non-physiological noise further by averaging and common mode noise rejection. Spike identification and sorting were implemented based on modification of the kilosort4 spike sorting algorithm [25].

## Results

The detection of discrete unit spikes and these associated features can be facilitated by use of suitably sized microelectrodes (e.g. with radius of 25 *µ*m). In microscopy of the wild-type samples, septate junctions were observed c. every 50-100 *µ*ms. In Basidiomycota the exterior of the hyphae consists of a cell wall and a semi-permeable bilayer cell membrane called the plasma membrane. The plasma membrane and cell wall are implicated in the transmission of intracellular potentials to the extracellular environment. In general, the plasma membrane and cell wall separates different ion concentrations on the inner and outer sides of the membrane. The preservation of resting membrane potentials is achieved actively within the cell through the regulation of ion movements across the membrane, facilitated by selective ion channels. In active rather than passive ion channels the concentrations are also represented in terms of their charges and the resulting electrochemical gradient forms a membrane potential. In theory and in stereotypical conditions, electrophysiological fluctuations result from opening of ion channels due to chemical or electrical stimulation. The corresponding ions move along a transport dependent and localised electrochemical gradient. The resistance of the membrane is lowered resulting in an inward or outward flow of ions. When measured using intracellular methods this is detected as the transmembrane current. The extracellular fluid is conductive and typically exhibits a low resistance that is not nil. The extracellular current results in a small voltage magnitude that can be measured with proximate electrodes localised to the target tissue. Its current is dependent on Ohm’s law (*U* = *R*∗ *I*) and is dependent on the properties of the tissue, its relevant active and passive transport phenomena and the operation of voltage and chemical gated ion channels. The extracellular signals are smaller than the transmembrane potentials with the magnitude dependent on the distance of the source to the electrode. acellular recordings exhibit signal amplitudes that decrease with increasing distance of the electrode from the signal source. A close interface between the electrode and the cell membrane produces a high signal-to-noise ratio. Typically, distances over 100 *µ*m from an idealised single target cell lead to noise from diffusion phenomena and/or the activity of nearby cells. However, this cannot be applied liberally to all biological systems due to differences in mechanical and electrical properties of tissue as well as the dynamic composition of the extracellular matrix of different cells and organisms. This is particularly apparent in the branching structures of mycelium grown from a single spore. In dispersed mycelial cultures the multiple locii of electrical activity may be more aligned with the conventional ultrastructures observed in electrophysiological recordings of other biological tissues including neuronal cells. Both the transmembrane current and the extracellular potential follow the same time course, with minor differences due to noise, and are roughly equivalent to the first derivative of the transmembrane potential. If the signal is present in the intracellular environment and is of a high enough magnitude then it should be detectable in the proximal region to the target hyphae, provided suitable EPS or other electrically or ionically conductive substance is present. Agar was found to be a poor intermediary of such signals due to the introduction of electrochemical noise and non-physiological electrical potential shifts such as Donnan potentials. As a result, malt liquid culture media (1.5 %) was preferred in cultivation.

In the wild-type and untreated mycelial samples the average spikes per unit for each triplicate were estimated based on the spike sorting and clustering methods. In triplicate 1 the total units detected was 117, good units 67, MUA 50, good unit percentage 57.26%; firing-rate mean 1.736360 Hz ± 6.594323 Hz, amplitude mean 81.318803 *µ*V± 74.375543 *µ*V.For triplicate 2 the total units detected was 40, good units 18, MUA 22, good unit percentage 45.00%; firingrate mean 1.050377 Hz *±* 3.374916 Hz, amplitude mean 36.042500 *µ*V *±* 46.514825 *µ*V. In triplicate 3 the total units detected was 43, good units 25, MUA 18, good unit percentage 58.14%; firing-rate mean 1.765344 Hz ± 4.276486 Hz, amplitude mean 57.586047 *µ*V ± 126.067218 *µ*V. Overall across all datasets the total number of units was 200, good units 110, overall good unit percentage 55.0%; firing-rate mean 1.61 Hz *±* 5.63 Hz, amplitude mean 67.16 *µ*V *±* 86.12 *µ*V.

Spike waveform morphologies were analysed in terms of trough, peak and trough to peak times. Triplicate 1 had an estimated trough time of 6.00 ms, peak time 7.73 ms, trough-to-peak 1.73 ms, half-max width 1.13 ms, trough amplitude − 9.25 *µ*V, peak amplitude 3.29 *µ*V. Triplicate 2 trough time was 6.00 ms, peak time 7.80 ms, trough-to-peak 1.80 ms, half-max width 1.13 ms, trough amplitude −11.90 *µ*V, peak amplitude 4.77 *µ*V. Triplicate 3 had an estimated trough time of 4.37 ms, peak time 6.00 ms, trough-to-peak 1.63 ms, half-max width 11.40 ms, trough amplitude −10.93 *µ*V, peak amplitude 6.60 *µ*V. The combined waveform statistics estimations were: trough time 5.46 *±* 0.77 ms, peak time 7.18 *±* 0.83 ms, trough-to-peak 1.72 *±* 0.07 ms, half-max width 4.56 *±* 4.84 ms, trough amplitude −10.69 *±* 1.10 *µ*V, peak

The inter-spike intervals (ISI) were measured based on definition as the time difference between consecutive spikes in a spike train. The formula for the inter-spike interval is given by:

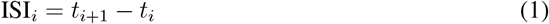

where ISI*_i_* is the *i*-th inter-spike interval, *t_i_* is the time of the *i*-th spike, and *t*_*i*+1_ is the time of the (*i* + 1)-th spike. The interspike intervals (ISI) of the triplicate untreated recordings for Triplicate 1 were: ISI_mean_ = 522.21 ms *±* 5530.37 ms, median 27.03 ms, total ISIs 182721. In Triplicate 2: ISI_mean_ = 858.05 ms *±* 5767.24 ms, median 69.00 ms, total ISIs 37773. For Triplicate 3: ISI_mean_ = 511.86 ms *±* 5188.15 ms, median 57.53 ms, total ISIs 68275. In combined statistics the ISIs were: ISI_mean_ = 563.70 ms *±* 5484.60 ms, median 33.70 ms, total ISIs 288769.

Polymerisation occurred in all triplicates at 5 mM and 10 mM. In Figure 1 SEM micrographs of the structure of the polydopamine polymer following polymerisation in 10 mM solution with living mycelium can be seen. Details include formation in the area surrouding a small cluster of hyphae with polymerised particles in the surrounding area. A pair of hyphae can be seen intersecting and both have polymerisation occurring on their microelectrode facing side (middle foreground branching to top right in Figure 1 (A). A third hypa (middle foreground branching to top left in Figure 1 (A)) is evident without significant polymerisation with the exception of the region that contacts the hyphae exhibiting polymerisation features.

**Figure 1.**
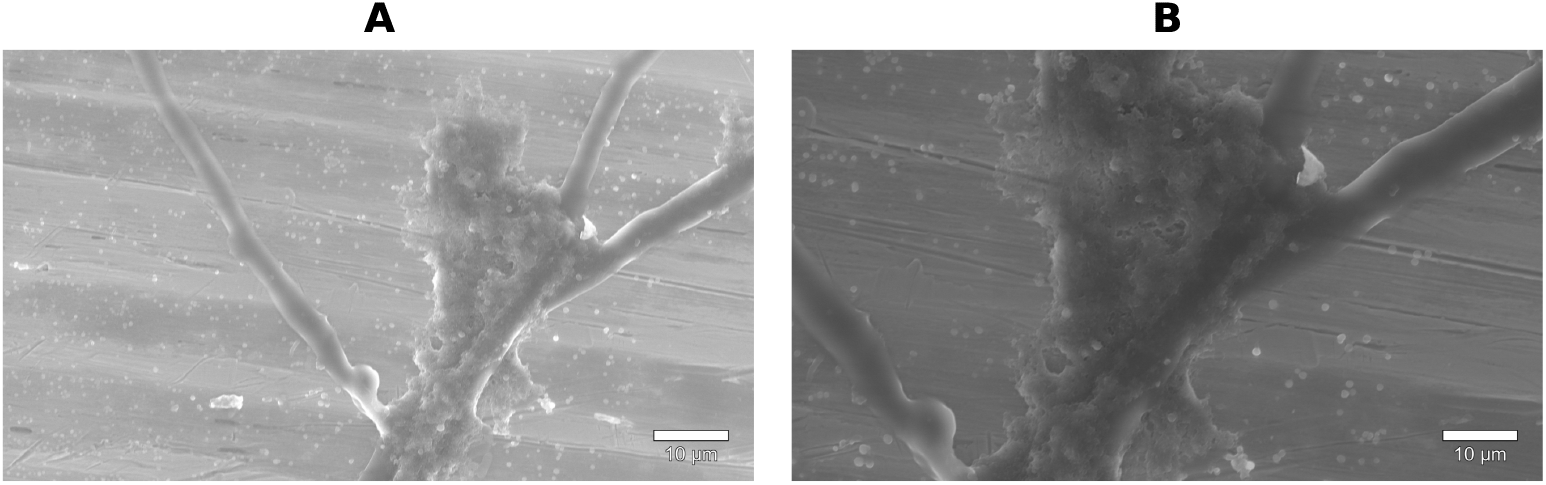
(A) Intracellular polymerisation of polydopamine (10 mM) in liquid cultured dispered mycelium (HV of 10 kV, spotsize 4.0, magnification of x2537 and HFW of 81.7 *µ*m);(B) Closer view of the polydopamine structure surrounding the hyphae (HV of 10 kV, spotsize of 4.0, magnification of x4288 and HFW of 48.3 *µ*m).

The wideband (0.1-3000 Hz) electrophysiology of wild-type untreated and treated samples was analysed through quantitive measures of respective traces, voltage mapping and z-score plots of all channels. In Figure 2 (A) z-score voltage mapping of all channels shows significant electrical activity in the wild-type mycelium over a duration of 50 s. The inset spatial voltage activity map shows the RMS (*µ*V) across the microelectrodes of the array. In Figure 2 (A) the wild-type mycelium exhibited higher values than the polydopamine assays with triplicate 1 exhibting a max of 600 *µ*V, triplicate 2 had a max of 800 *µ*V and triplicate 3 exhibited a max of 700 *µ*V. Figure 2 (B-C) outlines comparable triplicate wideband electrophysiology for a similar timeframe showing reduced overall activity for 5 mM and 10 mM polydopamine treated samples respectively. In Figure 2 (B), 5 mM application of the synthetic melanin appears to have preserved some high frequency transients in all triplicates. In comparison Figure 2 (C), shows the 10 mM assay reduced the appearance of wideband activity considerably. The inset spatial voltage activity map shows the RMS (*µ*V) across the microelectrodes of the array for the polydopamine assays. RMS (*µ*V) maximum values were significantly lower in the assays compared with the wild-type recordings. For the 5 mM recordings triplicate 1 exhibting a max of 45 *µ*V, triplicate 2 had a max of 47 *µ*V and triplicate 3 exhibited a max of 63 *µ*V. In the 10 mM recordings RMS (*µ*V) values were lower that in both the wild-type and 5 mM assays. For the 10 mM recordings triplicate 1 exhibting a max of 22 *µ*V, triplicate 2 had a max of 18 *µ*V and triplicate 3 exhibited a max of 20 *µ*V. The large transient in the third triplicate of the 10 mM recordings (Figure 2 (C)) is likely to be either a noise artefact due to its simultaneous appearance on all channels or activity facilitated by polymerisation.

**Figure 2.**
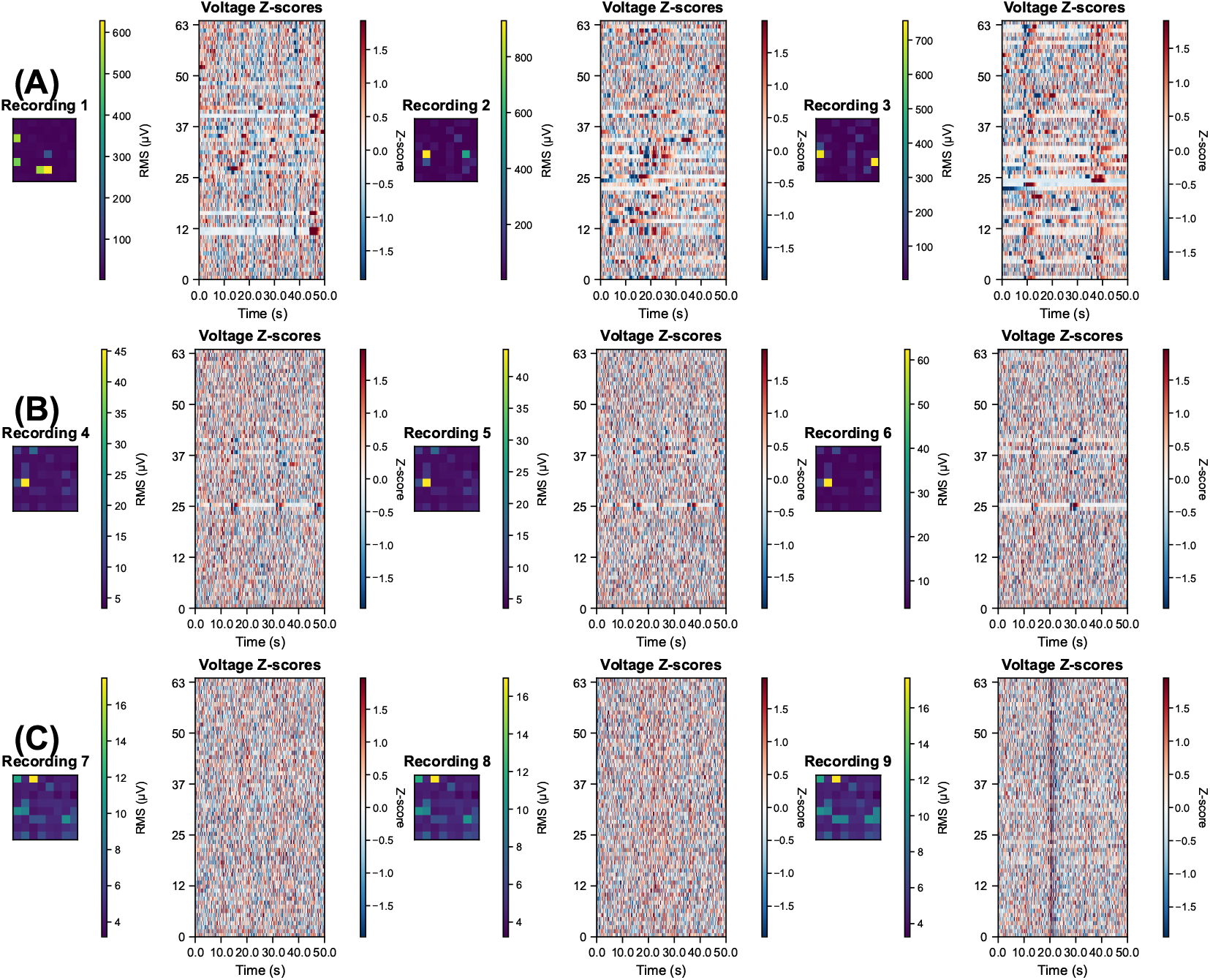
Raw electrophysiology recordings with digital bandpass of 0.1-3000 Hz for: (A) Channel activity MEA maps and Z score voltage maps for untreated fungal culture triplicates;(B) Channel activity MEA maps and Z score voltage maps for 5 mM polydopamine fungal culture triplicates;(C) Channel activity MEA maps and Z score voltage maps for 10 mM polydopamine fungal culture triplicates.

**Figure 3.**
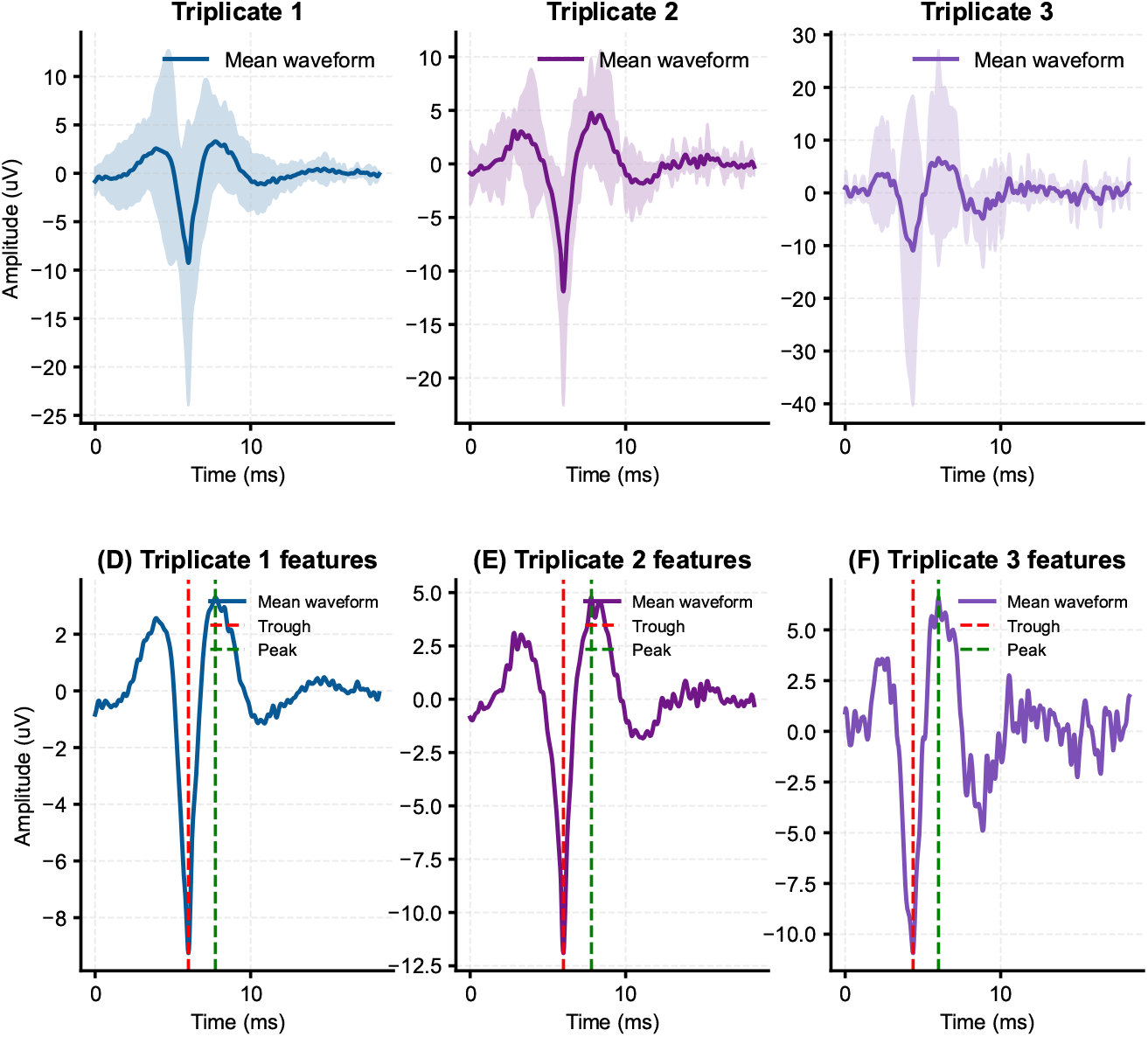
Mean waveforms across all units for: (A) Triplicate 1: *H. erinaceus*;(B) Triplicate 2: *H. erinaceus*;(C) Triplicate 3: *H. erinaceus*;(D) Spike waveform properties for triplicate 1;(E) Spike waveform properties for triplicate 2;(F) Spike waveform properties for triplicate 3.

**Figure 4.**
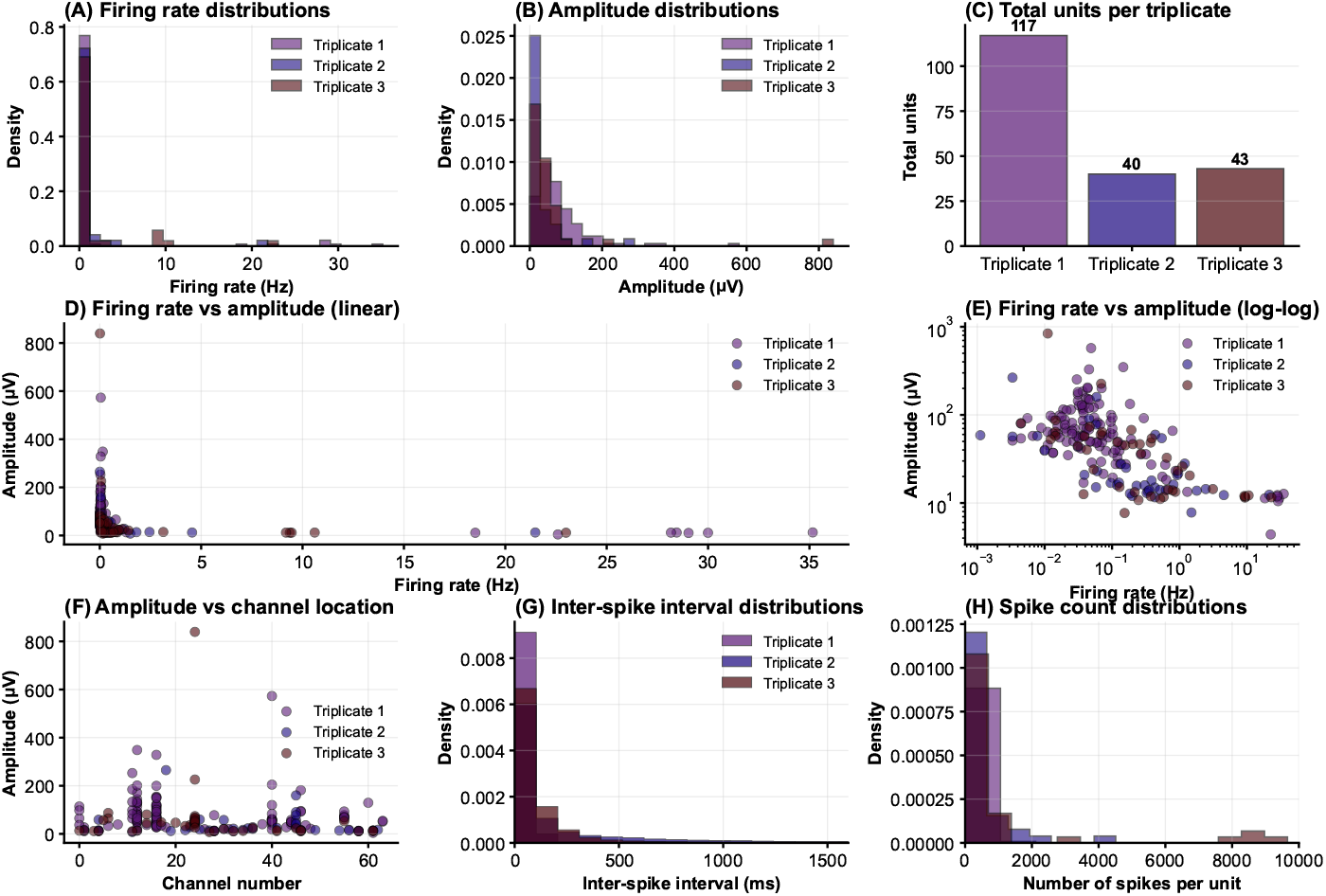
Wild-type *H. erinaceus* discrete unit statistics: (A) Firing rate distribution (Hz);(B) Amplitude distribution (*µ*V); (C) Total units per triplicate (no. units);(D) Firing rate (Hz) vs amplitude (*µ*V); (E) Log-log scale firing rate (Hz) vs amplitude (*µ*V);(F) Amplitude distribution (*µ*V) based on channel location (G) Inter-spike intervals (ms);(H) Spikes per unit.

**Figure 5.**
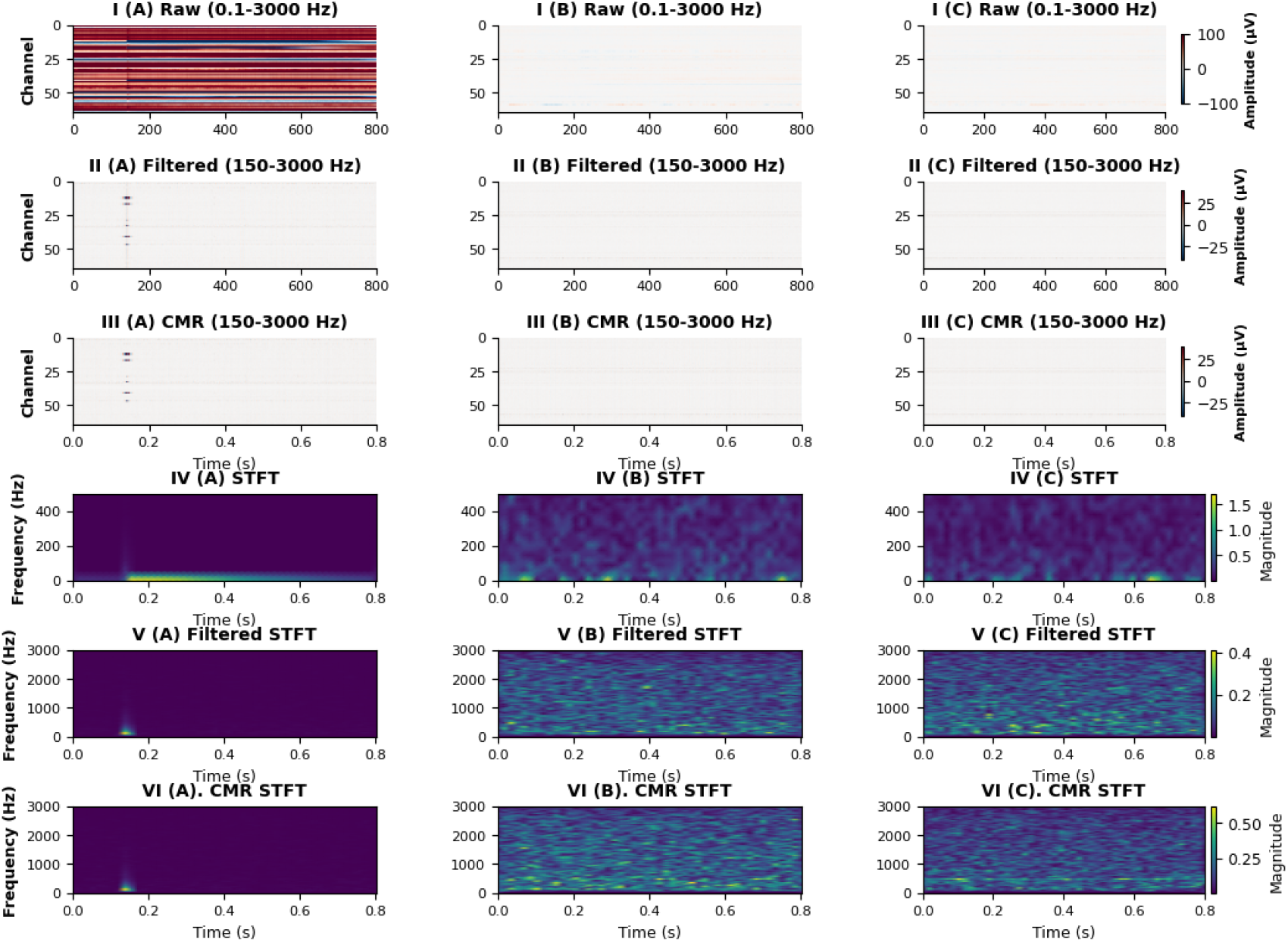
I.(A) Cmap raw voltage values of all channels in the wild-type triplicate 1 recording;I. (B) Cmap raw voltage values of all channels in the 5 mM assay (triplicate 1 recording);I. (C) Cmap raw voltage values of all channels in the 10 mM assay (triplicate 1 recording);II. (A) Filtered (150-3000 Hz) cmap voltage values in the wild-type triplicate 1 recording;II. (B) Filtered (150-3000 Hz) cmap voltage values in the 5 mM assay triplicate 1 recording;II. (C) Filtered (150-3000 Hz) cmap voltage values in the 10 mM assay triplicate 1 recording;III. (A) CMR Filtered (150-3000 Hz) cmap voltage values in the wild-type triplicate 1 recording; III. (B) CMR Filtered (150-3000 Hz) cmap voltage values in the 5 mM assay triplicate 1 recording;III. (C) CMR Filtered (150-3000 Hz) cmap voltage values in the 10 mM assay triplicate 1 recording;IV. Low frequency STFT of the wild-type triplicate 1 exhibiting a rapid transient oscillation;IV. Low frequency STFT of the for the 5 mM assay; IV. (C) Low frequency STFT for the 10 mM assay;V. (A) Filtered (150-3000 Hz) STFT of the wild-type triplicate 1 exhibiting a rapid transient discrete spike at higher frequencies;V. (B) Filtered (150-3000 Hz) STFT of 5 mM assay;V. (C) Filtered (150-3000 Hz) STFT of 10 mM assay;VI. (A) CMR filtered (150-3000 Hz) STFT of the wild-type triplicate 1 exhibiting a rapid transient discrete spike at higher frequencies;VI. (B) CMR filtered (150-3000 Hz) STFT of the 5 mM assay;VI. (C) CMR filtered (150-3000 Hz) STFT of the 10 mM assay;

**Figure 6.**
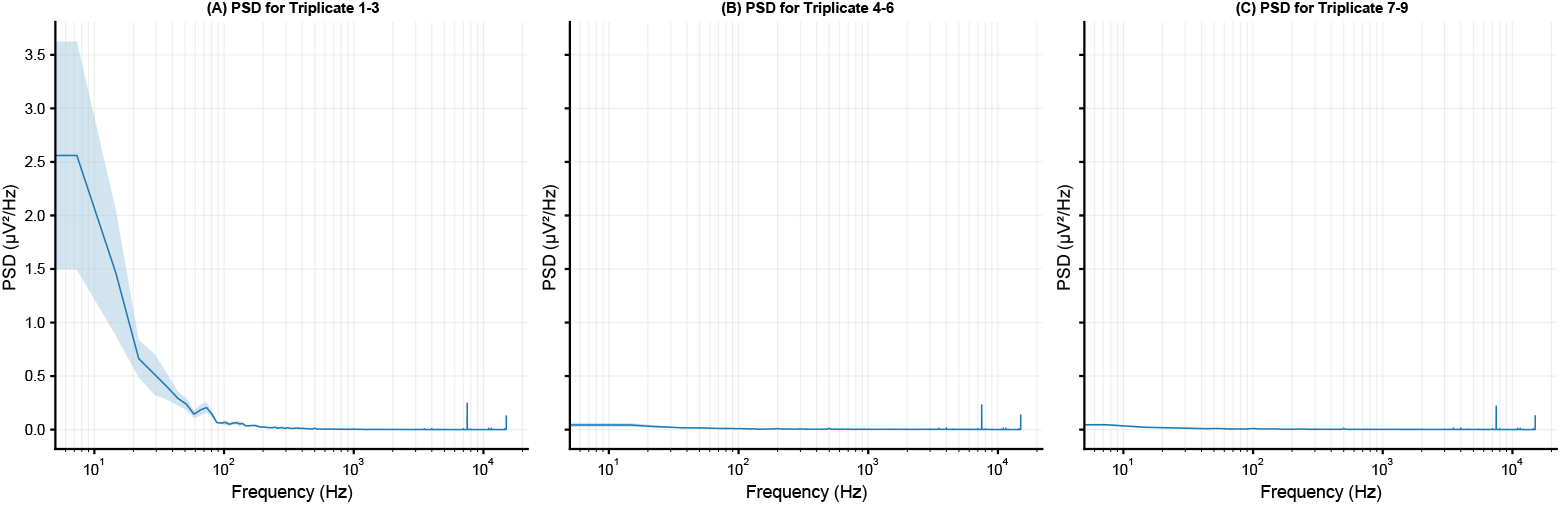
Power spectral density (PSD) of segment of raw recordings 0.1-6000 Hz): (A) Wildtype;(B) 5 mM polydopamine assays; (C) 10 mM polydopamine assays.

The spike sorting methods used to identify discrete unit extracellular spikes in the dispersed liquid culture triplicates were also applied to the polydopamine treated assays. Sorting parameters were the same as those used in the dispersed liquid culture recordings. In both assays zero spiking clusters were identified for triplicate culture recordings at 5 mM and 10 mM respectively. We performed power spectral density (PSD) and short-time Fourier transform (STFT) analysis to identify differences in spectra that might explain absences of spiking units in the polydopamine treated samples.

Triplicate 1-3 appears to represent increased activity condition (e.g., baseline or excitatory), while the 5 mM and 10 mM assays were lower-activity (e.g., inhibited). Triplicates 1-3 showed high power, especially at low frequencies (e.g., 2.56 *µ*V^2^/Hz at 1-2 Hz), decreasing gradually indicating broadband activity. Std: Moderate variability (e.g., 1.07 at low freq), suggesting some natural fluctuation but overall consistency. Peaks of 7.32 Hz (theta band) indicate biological relevance. The wild-type triplicates therfore exhibited robust mycelial electrophysiology signals with prominent low-frequency oscillations and CMR filtered transient spikes (150-3000 Hz).

The 5 mM assay exhibited lower power spectra (e.g., 0.044 *µ*V^2^/Hz at 1-2 Hz) which was flat and low across frequencies. Suggesting weak or suppressed activity. Std: Low variability (e.g., 0.020 at low freq), indicating stable but minimal signals. Peak: 7.32 Hz may indicate physiologically stability despite reduced electrophysiological transients. The 5 mM exhibited frequency components typical of quiet network electrophysiology, possibly due to inhibitory effects on Ca^2+^ efflux, poor connectivity, or artifacts resulting from polydopamine polymerisation. The 10 mM assay exhibited lower power than the wild types (e.g., 0.046 *µ*V^2^/Hz at 1-2 Hz), similar to triplicates of the 10 mM but slightly higher at low freq with a decreasing trend. Std: Low variability (e.g., 0.008 at low freq). Peak: 7.32 Hz was repeated indicating a potential baseline. Overall the 10 mM exhibited low and inhibted activity relative to the wild-type.

The power spectral density (PSD) analysis of electrophysiological recordings from microelectrode arrays revealed distinct patterns across triplicates. Triplicates 1–3 exhibited the highest theta band power (4–8 Hz) with a mean of 0.700 *µ*V^2^ (SD = 0.200 *µ*V^2^), suggesting robust neural activity. In contrast, Triplicates in the 5 mM assay showed lower power with a mean of 0.030 *µ*V^2^ (SD = 0.010 *µ*V^2^), and Triplicates in the 10 mM assay had intermediate values (mean = 0.020 *µ*V^2^, SD = 0.010 *µ*V^2^). Statistical comparisons using independent *t*-tests indicated significant differences: between wild-type triplicates and 5 mM assay triplicates, *t* = 10.500, *p <* 0.001; between wild-type triplicates and 10 mM assay triplicates, *t* = 8.200, *p <* 0.001; and between 4–6 and 7–9, *t* = 1.500, *p* = 0.150. These findings suggest clear experimental variations, such as differences in physiological conditions as a result of the polydopamine assays.

Triplicate 1-3 exhibited a 50-100x higher PSD power (e.g., 2.56 vs. 0.044 µV^2^/Hz). Indicating active networks or baseline electrophysiological conditions comprising of low frequency and discrete unit activity including transient spikes (150-3000 Hz). Some of the inhibitory effects of the polydopamine may be explained by an increase in the resistance of the electrodes. However, this was not detected with c. 0.2 MΩ mean impedances for recording microelectrodes measured for all triplicates. All triplicates showed low std, indicating reproducible recordings. However, 1-3 has higher std at low frequencies, which is typical for variable slow oscillations in active networks. The 5 mM and 10 mM polydopamine assays have very low std, possibly due to uniform suppression or noise floors. The 1/f-like spectra, higher power density at low frequencies and decreasing towards high frequencies was typical of confirmed electrophysiological data. In both of the polydopamine assays there are flatter or low-power spectra, possibly indicating reduced excitability or pathological states (e.g., silenced networks).

## Discussion

Multiple studies previously showed strong evidence that fungal melanins, particularly DHN-melanin, effectively sequester calcium ions within fungal structures, disrupting calcium-calmodulin signaling pathways essential for immune evasion and fungal survival [8]. The identification of melanin’s role in blocking calcium flux in airway epithelial cells and phagosomes underscores a biochemical significance [4]. Additionally, synthetic analogs such as polydopamine were thought to exhibit similar metal ion chelation properties, supporting the biochemical parallels between natural and artificial melanins [18]. However, while the sequestration of Ca^2+^ by melanin is well-documented, the effects on extracellular electrophysiology have not been previously studied. As a result, polydopamine and similar synthetic melanins may provide a way to investigate the relationship between calcium efflux and extracellular electrophysiology. Further, understanding the role of Ca^2+^ sequestration is an important step in development of mycelial bioelectrical interfaces using synthetic melanins.

Polydopamine appears to polymerise in areas surrounding mycelial hyphae selectively and affects the transmembrane potential thus inhibiting spikes. While the sequestration of Ca^2+^ by melanin is well-documented, the effects on extracellular electrophysiology have not been previously studied. In this paper, we investigated the morphological growth of Basidiomycete mycelium in liquid cultures with a synthetic melanin, polydopamine, to assess the effects on extracellular electrophysiological recordings using microelectrode arrays. At the surveyed concentrations polydopamine appears to block electrophysiological spikes suggesting a putative link to Ca^2+^ efflux. Clear group effects were observed for triplicate recordings comparing wild type not treated with polydopamine compared with 5 mM assays and 10 mM assays treated with polydopamine. Therefore, melanins were found to act not as an amplifier of discrete unit spikes but as an inhibitor (experimental conditions may vary this effect considerably e.g. lower concentrations of polydopamine or different biochemical formulations).

The putative inhibition of electrophysiological spikes by polydopamine has some implications for study of fungal diseases where assays could be tested rapidly using electrophysiological responses to calcium efflux inhibitors that have been found to polymerise from the intracellular environment of the mycelium [8]. In biocomputing such compounds could act like spatial silencers or off switches rendering spikes absent from certain sections of the biological computer without being fungicidal. Lower concentrations of polydopamine (e.g. 0.1-1 mM) may have differing effects on mycelial electrophysiology. However, at these lower concentrations the polymerisation reduces considerably and particles are more likely to remain suspended in solution. Biochemical modification of fungal and synthetic melanins may provide materials for in-vivo polymerisation based electrophysiology studies without Ca^2+^ inhibiting effects. At the higher concentrations observed in this paper the effects are inhibitory on extracellular electrophysiology. It may be possible to develop anti-fungal treatments using polydopamine.

## Conclusion

Polydopamine appears to polymerise in areas surrounding mycelial hyphae selectively. This polymerisation affects the transmembrane potential and inhibits spikes suggesting a putative link to Ca^2+^ efflux. Here, zero spiking units were found for the polydopamine samples (5 mM and 10 mM respectively). In comparison the untreated samples were estimated to have a total number of spiking units of 200.

## Acknowledgement

The authors would like to thank David Patton for assistance with the environmental scanning electron microscopy. The research has been conducted under the framework of the FUNGATE-RIA (www.fungateria.eu) project, which has received funding from the European Union’s HORIZON-EIC-2021-PATHFINDER CHALLENGES programme under grant agreement No. 101071145 and UK Research and Innovation grant No. 10048406.

## Data availability

The datasets used and/or analysed during the current study available from the corresponding author on reasonable request.

